# Abiotic pollen loss: The neglected pollen fate

**DOI:** 10.64898/2026.03.10.710827

**Authors:** Bruce Anderson, Sam McCarren, Ana Carolina Sabino-Oliveira, Vinicius L. G. Brito

## Abstract

**Background:** Pollen production is a costly investment in angiosperm reproduction, yet only a small fraction of grains reach conspecific stigmas. While pollen loss to floral visitors is well studied, the role of abiotic factors in shaping pollen fate has been largely overlooked. Understanding the relative contributions of biotic and abiotic pollen loss pathways is essential for interpreting floral trait evolution and pollination efficiency.

**Results:** We quantified abiotic pollen loss in four animal-pollinated species with contrasting floral longevities and reproductive phase dynamics. Flowers were monitored for five hours after anthesis, with pollen loss compared between unvisited flowers and those receiving single pollinator visits. Across all species, substantial pollen loss occurred in the absence of visitation, ranging from 37–57% of grains. In several cases, losses to legitimate pollinators were indistinguishable from abiotic loss alone, whereas pollen-foraging honeybees removed significantly greater fractions.

**Conclusion:** Abiotic factors can account for a large proportion of total pollen loss, sometimes equalling or exceeding pollinator-mediated removal. These findings challenge the assumption that pollen loss is primarily driven by pollinator activity and suggest that floral traits such as closure, gradual pollen release, and pollen packaging may function as adaptations that minimize environmental loss. Incorporating abiotic pollen loss into studies of pollen presentation and pollinator effectiveness provides a more complete understanding of selective pressures shaping floral evolution.

## Introduction

Pollen production is a costly, yet central component of angiosperm reproduction (Minnaar *et al.,* 2019). As the male gametophytes of flowering plants, each pollen grain represents a discrete unit of reproductive investment and a potential opportunity for a plant to sire offspring (Burd, 1994). However, because the number of pollen grains produced by a plant is limited, losses during transfer make the efficiency of pollen export and delivery a key determinant of the reproductive fitness (Westerkamp, 1997; Harder & Wilson 1998; Westerkamp & Claßen-Bockhoff, 2007). It is estimated that only small fractions (1-3%) of produced pollen reach the stigmatic surface of conspecific flowers (Harder & Thomson 1989; Holsinger & Thomson 1994; Harder *et al.,* 2000; Gong & Huang 2014; Johnson *et al.,* 2005). This potentially makes efficient pollen export one of the main challenges for fertility and an important area for natural selection to act upon (Minnaar & Anderson 2019). In fact, pollen grains embark on a particularly arduous journey as they move from their flower of origin to the bodies of pollinators and to the stigmas of recipient flowers. Along this journey, pollen is lost at multiple stages, and each stage of loss represents an opportunity for natural selection to act on traits that improve pollen transfer efficiency and, ultimately, the proportion of pollen grains that achieve successful fertilization (Anderson & Minnaar 2020). Classic theories such as the “pollen economy” (Lloyd 1984) and the “pollen presentation” theory (Harder & Thomson 1989) emphasize the importance of efficient pollen use, yet it remains unclear which stages along the pollen journey are responsible for the greatest sources of pollen loss and, consequently, where selection on pollen transfer efficiency is likely to be strongest.

Pollen loss caused by floral visitors that directly consume pollen or collect it for their nests is well documented in the literature. For example, substantial amounts of pollen are removed by bees, which groom pollen from their bodies and actively pack it into their scopae or corbiculae (Thomson, 1986; Thorp, 2000; Portman *et al.,* 2019). This pollen is transported to the nest, where it represents the primary source of protein for larval development (Harder, 1990; Hargreaves *et al.,* 2009). In addition to bees, pollen is directly consumed by a wide range of pollen-feeding floral visitors, including flies (Holloway, 1976; Haslett, 1989; Brodie *et al.,* 2015), beetles (Johnson & Nicolson 2001), micropterigid moths (van der Pijl, 1960), heliconiid butterflies (Gilbert, 1972), and some nectar-feeding bats that supplement their diet with pollen (Herrera & Martínez del Río, 1998). In these cases, pollen is removed from the pollination pathway by floral visitors, contributing directly to pollen loss. Large pollen fractions can also be lost during foraging when pollinators dislodge pollen, causing it to fall from anthers or stigmas onto the ground or petals (Harder & Thomson, 1989). Furthermore, pollen grains from different flowers can interfere with one another on the bodies of pollinators so that pre-existing pollen grains on the pollinator prevent new pollen grains from attaching (Moir & Anderson, 2023; Brito *et al., in review*), or pollen grains from newly visited flowers smother, displace or remove pollen grains which already reside on the pollinator (Anderson *et al.,* 2024; Santana *et al.,* 2025). Although potentially substantial, these sources of pollen loss only occur during the brief intervals of pollinator visitation or, affect only the small number of pollen grains that actually attach to pollinators.

However, pollen may also be lost before pollinators even interact with the flower and this loss potentially acts on all of the pollen residing within a flower. For example, adverse abiotic conditions can damage or remove pollen from flowers prior to its placement on pollinators, potentially imposing selective pressures on floral traits that reduce pollen exposure to the abiotic environment. When Reynolds *et al.,* (2009) excluded pollinators from *Silene* flowers at night, they found that the flowers still lost up to 50% of their pollen grains, suggesting that wind may strip pollen from anthers or that pollen may be lost passively to the abiotic environment. Traits that may reduce abiotic pollen loss may include floral orientation to reduce water damage (Aizen, 2003; Huang *et al.,* 2002; Shibata *et al.,* 2025); nocturnal floral closure to prevent pollen from being exposed to nighttime moisture (Von Hase *et al.,* 2006); and anther closure to prevent rain damage (Humphreys & Skema, 2023). Corolla closure, which restricts potential pollinator access has also been shown to reduce pollen loss by up to 59% compared to open flowers (Bynum & Smith, 2001). Although some traits (e.g. nocturnal flower closure) potentially protect pollen when pollinators are inactive, other traits (e.g. gradual pollen release from anthers) potentially reduce abiotic pollen loss while pollinators are active, potentially resulting in an interesting trade-off in selective pressures to optimize pollen export: traits that protect pollen from abiotic loss may also reduce pollinator visitation or the amount of pollen that is transferred to pollinators. While we often view floral traits as adaptations that promote efficient pollen transfer between flowers and pollinators, we seldom interpret floral adaptations considering their potential to limit abiotic pollen loss. Naturally, the biotic and the abiotic routes to pollen loss are closely intertwined because the less pollen removed by pollinators, the more susceptible it is to abiotic pollen loss.

The susceptibility to pollen loss is likely also dependent on floral morphology. For example, differences in anther exposure, flower orientation, or the degree of anther dehiscence can influence how much pollen is effectively retained until a pollinator arrives (Buchmann, 1983; Harder & Thomson, 1989; Ushimaru et al. 2007). Thus, different floral architectures may modulate the balance between biotic and abiotic pollen losses, ultimately influencing plant reproductive efficiency. Given the potential importance of pollen loss before and between pollinator visits and the fact that few studies have directly quantified abiotic pollen losses, our study aimed to quantify the abiotic pollen loss in animal-pollinated species and to compare it with pollen removal by floral visitors. We used plant species that are visited by different pollinator species and distributed across two different continents. We hypothesized that even in the absence of pollinator visits, flowers may lose significant proportions of their pollen, perhaps even equivalent to the amount of pollen removed by the floral visitors themselves

## Methods

This study focused on four species with contrasting floral longevities and sexual phase dynamics: *Jacquemontia sp.* Convolvulaceae (flowers last a single day)*, Pyrostegia venusta* Bignoniaceae (flowers last two days, with the first day in the pollen-presenting phase and the second in the pollen-receiving phase)*, Pelargonium quercifolium* Geraniaceae (pollen-dispersing phase lasts approximately 1 day and anthers are shed on day 2) and *Lapeirousia anceps* Iridaceae (flower persist for 2-5 days, with anther dehiscence occurring on the first day and stigma unfurling on the second day) (Fig. 1). These species were selected to encompass a range of floral longevities, reproductive phase dynamics, and exposure times of pollen to both abiotic conditions and floral visitors.

**Figure 1:**
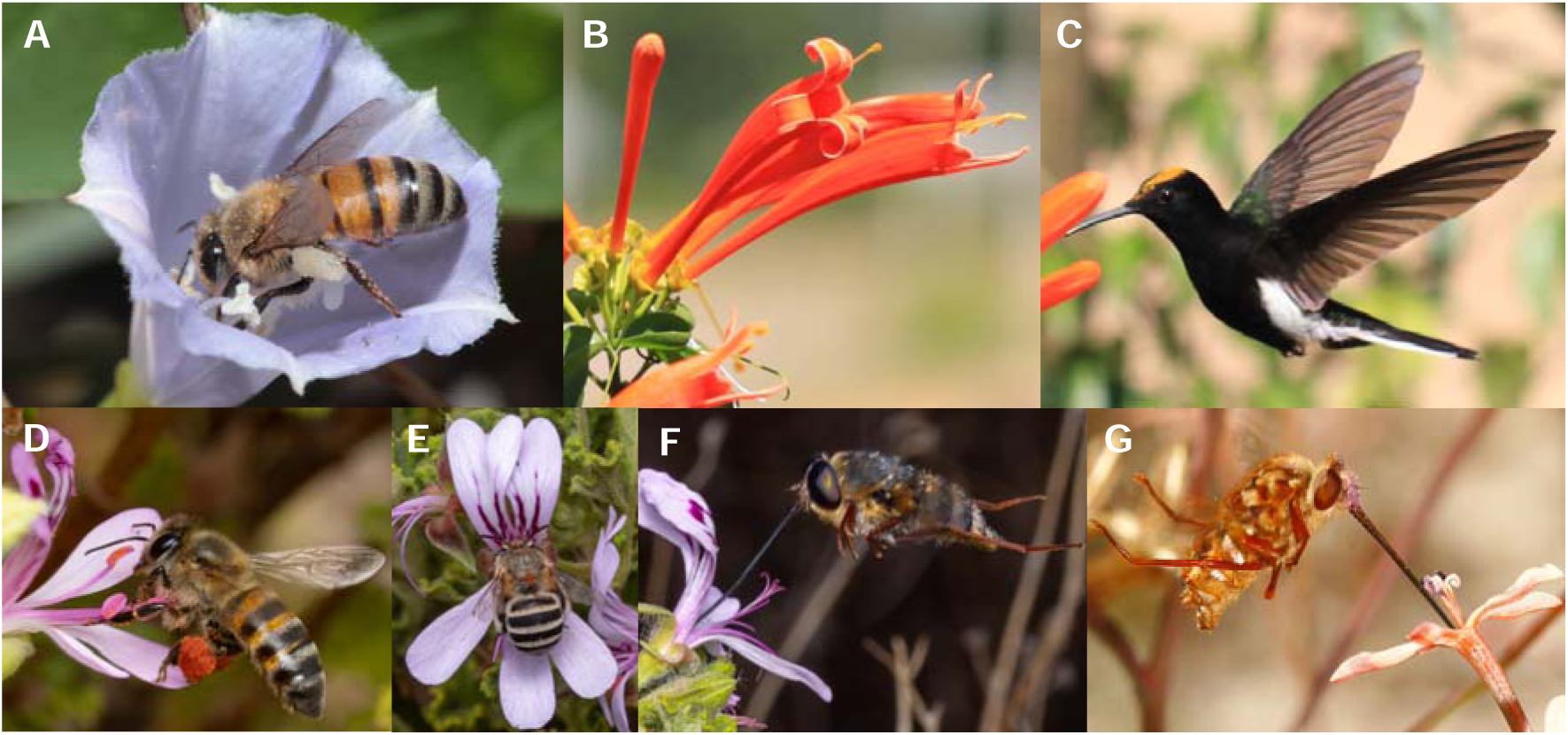
Flowers and their pollinators of four different systems used to quantify abiotic pollen loss in this study. A) *Jacquemontia sp.* flower being visited by a honeybee which harvests pollen directly from the anthers, without foraging for nectar. B) Long, tubular flowers of *Pyrostegia venusta.* C) The Black Jacobin hummingbird (*Florisuga fusca*) has a clear pollen load on its head after visiting *P. venusta flowers*. D) A honeybee collecting pollen from *Pelargonium quercifolium*. Honeybees do not forage for nectar but use their forelegs to actively remove pollen from the anthers. This is then transferred to the scopae on their hind legs. E) *Amegilla sp.* bee foraging for nectar in *P. quercifolium*, which place pollen on the ventral surface of the bee. F) Long-proboscid fly (*Prosoeca longipennis*) foraging for nectar in *P. quercifolium*. These flies seldom make contact with the reproductive parts of the flower and were never seen carrying large pollen loads from *P. quercifolium.* G) The long-probloscid fly, *Moegistorynchus longirostris* visits the long-tubed flower, *Lapierousia anceps*, which places its purple pollen on the forehead of the fly. Photographs D,E and F by Grant Evans.

Flowers of *Jacquemontia sp.* and *P. venusta* were sampled in Brazil near the JK Campus of the Federal University of the Jequitinhonha and Mucuri Valleys (UFVJM), in Diamantina, Minas Gerais, Brazil (−18.1667, -43.5000) while *P. quercifolium* and *L. anceps* were sampled in South Africa in the Baviaanskloof, Eastern Cape (−33.4912, 23.6038) and Mamre, Western Cape (−33.5214, 18.4545) respectively. We marked a total of 42 *Jacquemontia sp.* flowers, 36 *P. venusta* flowers, 59 *P. quercifolium* flowers and 100 *L. anceps* flowers. From each marked flower, one anther was removed in the morning just after anthesis to estimate total pollen production by an anther. Then, the flowers were observed for five hours, and any visitors were recorded. After five hours of observation, a second anther was collected so that pollen in this anther could be compared with pollen found in the first anther, the difference representing pollen loss after five hours. Pollen lost from flowers that remained unvisited after five hours was attributed to the abiotic environment. To ensure that some flowers remained unvisited, we occasionally had to keep pollinators away from flowers by chasing them off plants.

Collected anthers were stored in Eppendorf tubes with ethanol (200µl for *Jacquemontia sp.* and *P. venusta* and 1000µl for *P. quercifolium* and *L. anceps*). Later, the pollen grains in a subsample (10µl for and *L. anceps* and 15µl for *P. quercifolium*) from each tube were counted under the compound microscope. For the counts of *Jacquemontia sp.* and *P. venusta* we used a subsample of 10µl and a haemocytometer to assist with counting. The counts were further extrapolated to estimate the total number of pollen grains in each sample.

All statistical analyses were carried out in R (R Core Team, 2020). The pollen counts for all four species were scaled by dividing them by the mean of pollen production for each species to make them more easily comparable to each other. Then, using paired t-tests or for non-normal data a paired Wilcoxon signed rank test, we compared pollen production in each species to pollen remaining after receiving no visits. Further, using unpaired one-sided t-tests or an unpaired one-sided Wilcoxon signed rank test, we compared pollen loss (production – remaining) after receiving no visits and after receiving one visit. We also compared pollen loss after a single visit from a honeybee to *P. venusta* to pollen loss after no visits using a one-sample Wilcoxon signed rank test.

Additionally, a general linear model with quasipoisson distribution was used to compare pollen remaining in response to the number of visits for the species *Jacquemontia sp.* and *P. quercifolium,* which received up to 15 visits. The two other species were not included in this model because they received a maximum of two visits.

## Results

### Pollinator observations

*Jacquemontia sp.* was mainly visited by non-native honeybees (*Apis mellifera*). These bees were primarily collecting pollen as they seldom entered the depths of the flower in search of nectar and their scopae contained large quantities of *Jacquemontia sp.* pollen (Fig. 1A). The bodies of the bees had sparsely scattered *Jacquemontia* sp. pollen grains. *P. venusta* was visited by long-billed, nectar-feeding hummingbirds (Fig. 1B,C), including Swallow-tailed hummingbirds (*Eupetomena macroura*) and Jacobins hummingbird (*Florisuga sp.*). The birds had very clear, dense pollen loads on their foreheads. In contrast, there was a single visit from a non-native honeybee (*Apis mellifera*) that was unable to access nectar from the long corollas but harvested pollen from the anthers. *Pelargonium quercifolium* had a diverse assemblage of visitors, including native honeybees, which collected pollen straight from the anthers without attempting to access the nectar (Fig 1D). These bees had dense orange *P. quercifolium* pollen loads on their scopae and sparsely distributed pollen grains on the ventral sides of their bodies. *Amegilla sp.* foraged for nectar (Fig 1E) and had clear pollen loads on their ventral surfaces, while long-proboscid flies *Prosoeca ganglbauri* and *Prosoeca longipennis* foraged for nectar without making clear contact with the reproductive parts of the plant (Fig 1F). Long proboscid flies did not have clear *P. quercifolium* pollen loads that were visible with the naked eye. These insects are expected to primarily be nectar robbers. *Lapeirousia anceps* was mainly visited by the long-proboscid fly *Moegistorhynchus longirostris* (Fig 1G), which is known as its pollinator (Anderson *et al.,* 2016). They had dense loads of purple pollen on their frons. We also observed a few visits by a small bombyliid and a tiny solitary bee, which collected pollen without foraging for nectar. These insects did not come to our experimental flowers.

### Pollen loss

When the flowers received no visits after five hours of being exposed to the abiotic environment, a significant amount of pollen was lost from the anthers of all four species (Fig. 2). An average of 37.1% of pollen was lost from *Jacquemontia sp.* flowers (t = 4.22, df = 14, p < 0.001), 56.9% was lost from *P. venusta* flowers (V = 210, p < 0.001), 43.0% was lost from *P. quercifolium* flowers (t = 3.09, df = 20, p = 0.003) and 47.0% was lost from *L. anceps* flowers (t = 7.12, df = 73, p < 0.001).

**Figure 2:**
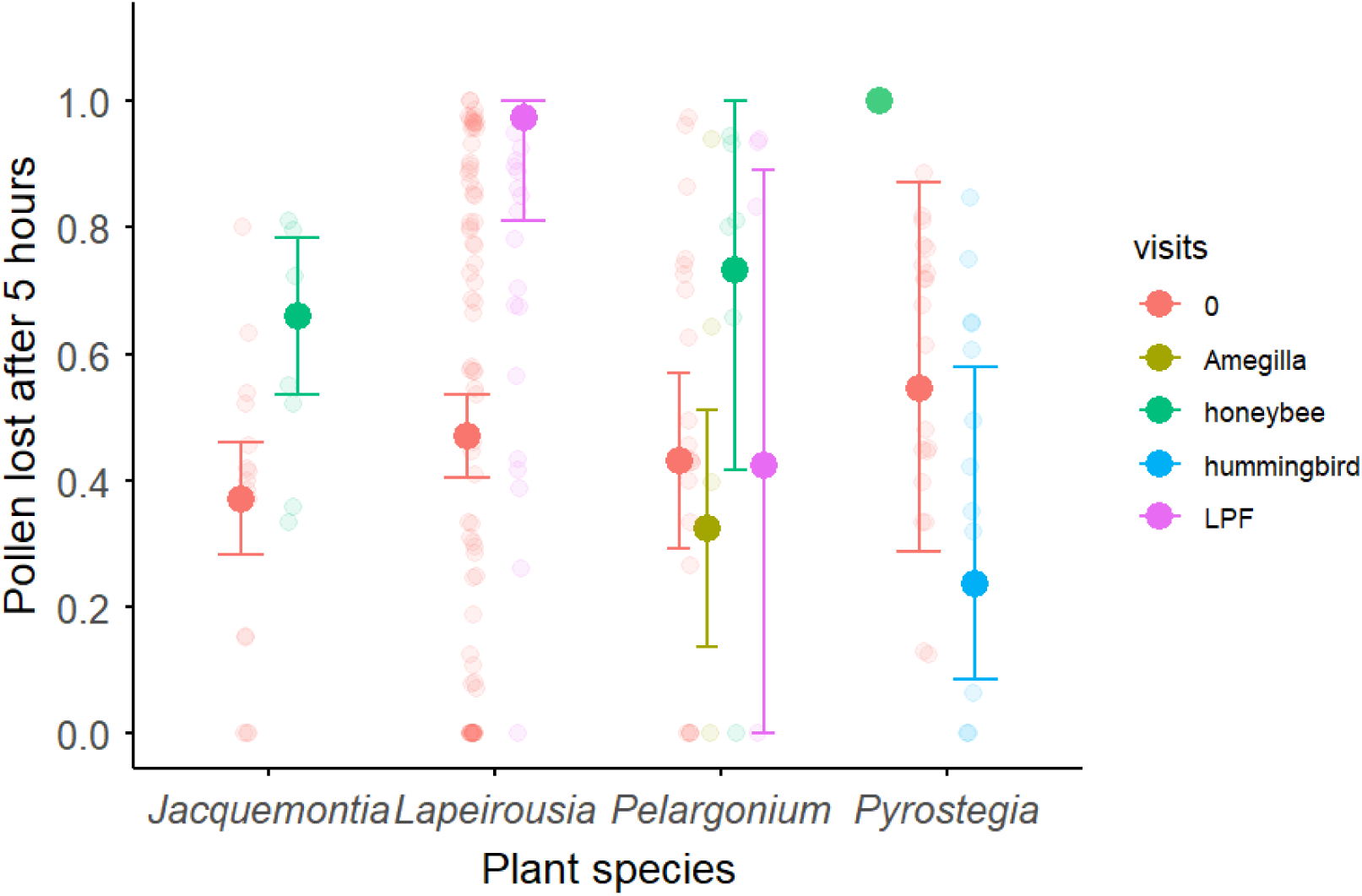
Standardised pollen loss from the anthers of Jacquemontia sp., L. anceps, P. quercifolium and P. venusta. five hours after anthesis when they had received no visits or one visit. Bold dots indicate the mean for Jacquemontia sp., L. anceps and P. quercifolium, the median for no visits and hummingbirds in P. venusta, and one data point for honeybees in P. venusta. Error bars indicate the standard deviation for Jacquemontia sp., L. anceps and P. quercifolium, and quartiles for P. venusta. Transparent dots represent the raw data.

In *Jacquemontia sp.,* visitation by a single honeybee plus potential abiotic pollen loss after five hours accounted for a 66.0% loss of pollen, which is significantly more than 37.1% attributed to the abiotic environment alone (t = 1.87, df = 20, p = 0.038). In *P. venusta,* loss attributed to a hummingbird visitor plus the abiotic environment did not remove more pollen than the abiotic environment alone (W = 77.5, p = 0.953), nor did single visits to *P. quercifolium* (t = 0.42, df = 33, p = 0.338). However, the single visit by a honeybee to *P. venusta* together with the abiotic environment removed significantly more pollen than the abiotic environment alone (V = 13, p < 0.001). Likewise, in *L. anceps* long-proboscid flies and the abiotic environment together removed about 94.8% of pollen, which is significantly more than the abiotic environment alone (t = 3.28, df = 93, p < 0.001).

The model testing pollen loss in response to species and visit number showed no effect of species by itself (Estimate = -0.22, LJ^2^ = 0.10, p = 0.756), but as the number of visits increased more pollen was lost (Estimate = -0.52, LJ^2^ = 23.55, p < 0.001, Fig. 3). Also, there was a significant interaction between plant species and number of visits (Estimate = 0.38, LJ^2^ = 6.32, p = 0.012), indicating that the rate at which pollen is lost across successive visits differs among species.

**Figure 3:**
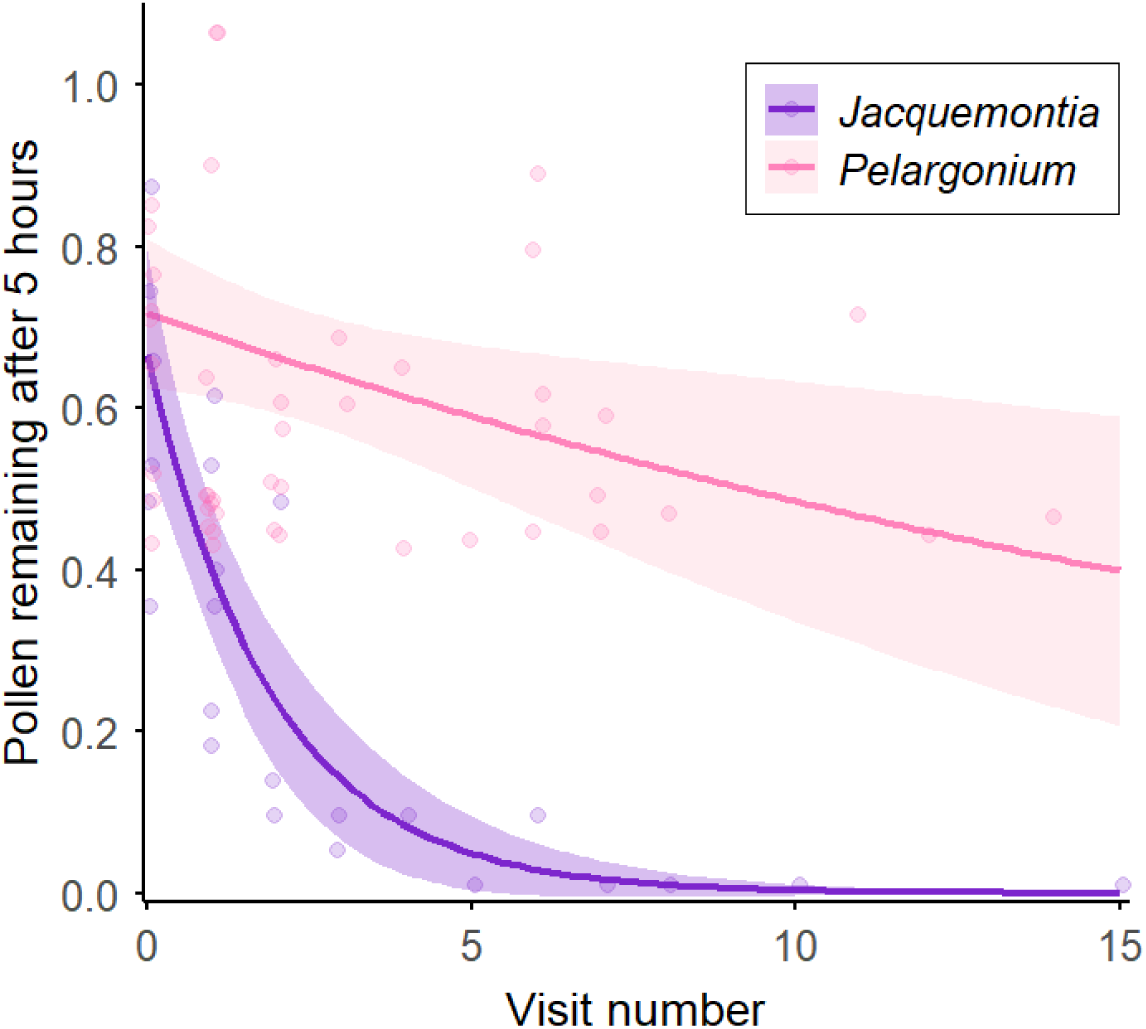
Pollen remaining in the anthers of *Jacquemontia sp.* and *Pelargonium quercifolium* five hours after anthesis in relation to number of visits as a continuous variable. Shaded areas indicate CIs.

## Discussion

In all studied species, pollen loss attributed to abiotic factors alone (i.e. flowers that were never visited) was substantial (37-56%) during the first five hours after anthesis. In contrast, pollen removal following single visits by legitimate pollinators was often indistinguishable from losses observed in unvisited flowers, suggesting that legitimate pollinators may often remove only small fractions of a floweŕs pollen load with each visit. In comparison, pollen removal by pollen-foraging honeybees was frequently much greater than abiotic pollen loss alone. Together, these results indicate that abiotic factors can account for a considerable proportion of total pollen loss and may even exceed the pollen removal resulting from pollinator interactions. Therefore, pollen loss cannot be assumed to result solely from pollinator activity, as abiotic losses may equal or surpass those caused by floral visitors. If this is the case, selection for floral traits that reduce abiotic pollen wastage may be as strong as, or even stronger than, selection imposed by pollinators to reduce pollen wastage. High levels of abiotic pollen loss also have important methodological implications: quantifying pollinator-mediated pollen removal requires explicit controls for abiotic loss and direct comparisons with unvisited flowers. Below, we examine pollen loss to floral visitors and to the abiotic environment in greater detail.

Hummingbirds did not remove detectable pollen loads from *P. venusta* flowers, nor did *Amegilla sp.* bees from *Pelargonium quercifolium*, suggesting that pollen transfer to legitimate pollinators may often account for only a small fraction of a flower’s pollen. This may seem contradictory to the observation that non-grooming pollinators often carry conspicuous pollen loads on their bodies (e.g. Moir *et al.,* 2022, Fig 1C and G in this paper). However, pollen loads are not necessarily indicative of how much pollen is transferred per visit, as pollen loads on non-grooming pollinators may accumulate over visits to many previous flowers (Price & Waser 1982). Indeed, studies with quantum dots show that non-grooming pollinators may become saturated with pollen so that flowers are unable to place significant quantities of pollen onto pollinators when they do visit (Moir & Anderson 2023; Brito *et al.,* in review).

The only legitimate pollinator in our study to remove quantifiable amounts of pollen was the long-proboscid fly visiting *L. anceps* (Fig. 1G). This fly appeared to remove nearly half of a flower’s pollen during a single visit, a loss that could not be explained by abiotic factors alone. In contrast, Minnaar et al. (2019), at the same study site used quantum dots to show that *L. anceps* flowers actually deposit an average of only 13.2 pollen grains per visit onto their long-proboscid fly pollinators. Together, these results highlight a striking discrepancy between pollen removal and effective pollen placement. This discrepancy may be explained if large quantities of pollen are dislodged during visits and are lost when they fall to the ground rather than being retained on the pollinator’s body. Such losses may be particularly pronounced when pollinators arrive already saturated with pollen and additional grains cannot adhere effectively (Moir & Anderson 2023; Brito *et al.,* in review), increasing the likelihood that newly dislodged pollen is shed rather than transported (see Fig 2b in Minnaar *et al.,* 2019). Harder and Thomson (1989) found that honeybees and bumblebees dislodged approximately 14% of a flower’s available pollen per visit while actively collecting pollen. However, honeybees and bumblebees groom frequently and so comparable or even greater dislodging may occur when pollinators arrive with saturated pollen loads, as limited adhesion space could increase passive shedding rather than retention. In contrast, we detected no discernible pollen removal by long-proboscid flies visiting *P. quercifolium*. The proboscides of these flies greatly exceeded the length of the floral tubes, and photographs indicate minimal contact with the reproductive structures (Fig. 1F), suggesting that these flies function primarily as nectar thieves rather than legitimate pollen vectors of *P. quercifolium*.

Honeybees were also observed visiting *P. quercifolium*, but they primarily foraged for pollen rather than for nectar (Fig. 1D). Although these bees appeared to remove substantial amounts of pollen, high variation among visits meant that pollen removal following single visits did not differ significantly from abiotic pollen loss alone. In contrast, single honeybee visits to *Jacquemontia sp.,* which is not pollinated by honeybees in its native habitat, resulted in substantial pollen removal per visit (Fig. 1A) and after five honeybee visits, there was almost no pollen remaining in the anthers (Fig. 3). Similarly, the individual honeybee visit to *P. venusta* also resulted in very high pollen loss. Overall, these results suggest that legitimate, non-pollen foraging pollinators may remove only small fractions of per visit, whereas pollen-foraging insects can extract large proportions of the available pollen. Consistent with this pattern, previous studies have reported that bees may remove more than 80% of a flower’s pollen in a single visit when they are actively foraging for pollen (Dunham, 1939, Strickler, 1979, Harder & Thomson 1989).

While studies on abiotic pollen loss are in their infancy, it is important to acknowledge that abiotic pollen loss may be important and, in some cases, may exceed pollen transfer and loss to pollinators. Recognizing the importance of abiotic pollen loss offers a new perspective for interpreting many floral traits. For example, adaptations commonly perceived as a response to pollinator-mediated selection may also function to reduce pollen exposure to abiotic stressors and limit passive pollen loss.

Such adaptations may be especially favored in long-lived flowers, where prolonged exposure to abiotic conditions increases the risk of cumulative abiotic pollen loss. If long-lived flowers experience greater cumulative exposure to abiotic stress, they may be under stronger selection to minimize pollen loss. Such selection could favour increasingly refined pollen protection and release mechanisms. These may include packaged pollen, as in orchids and asclepiads (Lloyd & Yates 1982), as well as gradual pollen release through poricidal or slit-like anthers, (Brito *et al.,* 2021; Vasques-Castro *et al.,* 2025), or the retention of pollen within protective floral structures until it is released in a single event (e.g., explosive pollination; Anderson et al. 2024). Traditionally, variation in pollen release strategies has been interpreted through the lens of pollen presentation theory (Harder & Thomson 1989), which predicts that gradual pollen release is favoured under high visitation rates, whereas rapid or complete pollen release is advantageous when visitation is infrequent.

However, our results raise the possibility that some forms of pollen packaging and release may also function to limit abiotic pollen loss. Under this view, both slow-release systems (e.g., poricidal anthers) and rapid-release systems (e.g., orchid pollinaria) could represent alternative solutions to the same problem: protecting pollen from environmental loss until effective transfer occurs. Disentangling the relative contributions of pollinator-mediated and abiotic selection in shaping pollen presentation strategies represents a promising direction for future research.

One potential caveat to our study is that the biological significance of pollen loss measured five hours after anthesis likely depends on floral longevity. In short-lived flowers such as *P. venusta* and *Jacquemontia sp.*, five hours may encompass much of the effective pollen-dispersing phase and so pollen loss that occurs mostly at the end of this period may not have evolutionary consequences. In contrast, in longer-lived flowers such as *P. quercifolium* and *L. anceps*, substantial dispersal opportunities might remain beyond this time frame. This raises the question of whether abiotic pollen loss occurs steadily throughout the pollen-presenting phase or whether it accelerates toward the end of the floral lifespan. Understanding the temporal dynamics of pollen loss across the lifespan of a flower would help clarify the overall impact of abiotic pollen loss on reproductive success.

Importantly, our results demonstrate that pollen loss from flowers cannot be assumed to result solely from pollinator activity. Substantial proportions of pollen may be lost to abiotic factors before or independently of visitation, with important consequences for how we estimate pollen transfer efficiency and pollinator effectiveness. Accurate assessments of pollen removal and transfer therefore require explicit controls for flower age and exposure time, as well as comparisons with unvisited flowers. More broadly, incorporating abiotic pollen loss into studies of pollen presentation and pollinator effectiveness may lead to a more complete understanding of the selective forces shaping floral evolution.

## Acknowledgements

We wish to thank Mark Trout and Vaalwater Lodge for permission to work in South Africa on *Lapeirousia anceps* and *Pelargonium quercifolium* respectively. Permission to work in Brazil was acquired through the licence SISBIO 53417. We also wish to thank Jason Cane, Nerine van Wyk and Kelvin Francis for help with field work and to Grant Evans for allowing us to use his photographs. BA would like to thank the Oppenheimer Memorial Trust for financial support.

## Notes

### Competing Interest Statement

The authors have declared no competing interest.

## References

Aizen MA. 2003. Down-facing flowers, hummingbirds and rain. Taxon 52: 675–680. *America* 69, 1403–1407.

Anderson B, Minnaar C. 2020. Illuminating the incredible journey of pollen. American Journey of Botany 107: 1–4.

Anderson B, Sabino-Oliveira AC, Matallana-Puerto CA, Arvelos CA, Novaes CS, Calaça DCC, Schulze-Albuquerque I, Pereira JPS, Borges JO, de Melo LRF, Consorte PM, Medina-Benavides S, Andrade TO, Monteiro TR, Marcelo VG, Silva VHD, Oliveira PE, de Brito VLG. 2024. Pollen wars: Explosive pollination removes pollen deposited from previously visited flowers. The American Naturalist 204: 616–625.

Anderson B, Pauw A, Cole WW, Barrett SC. 2016. Pollination, mating and reproductive fitness in a plant population with bimodal floral-tube length. Journal of Evolutionary Biology 29: 1631–1642.

Brito, et al. Pollen landscape saturation reduces pollen deposition by subsequently visited bat-pollinated flowers. Manuscript in review.

Brodie BS, Smith MA, Lawrence J, Gries G. 2015. Effects of floral scent, color and pollen on foraging decisions and oocyte development of common green bottle flies. PLoS One 10: e0145055.

Buchmann SL. 1983. Buzz pollination in angiosperms. In: Jones CE, Little RJ, eds. Handbook of Experimental Pollination Biology. New York, NY, USA: Scientific and Academic Editions, 73–113.

Burd M. 1994. Bateman’s principle and plant reproduction: the role of pollen limitation in fruit and seed set. The Botanical Review 60: 83–139.

Bynum MR, Smith WK. 2001. Floral movements in response to thunderstorms improve reproductive effort in the alpine species *Gentiana algida* (Gentianaceae). American Journal of Botany 88: 1088–1095.

De Brito VLG, Leite FB, Jorge LR, Sazima M. 2021. Distinct pollen release dynamics between stamens generate division of labour in pollen flowers of two *Pleroma* species (Melastomataceae). Flora 285:151961.

Dunham WE. 1939. Collecting red clover pollen by honey bees. Journal of Economic Entomology 32: 668–670.

Gilbert LE. 1972. Pollen feeding and reproductive biology of *Heliconius butterflies*. Proceedings of the National Academy of Sciences of the United States of

Gong YB, Huang SQ. 2014. Interspecific variation in pollen–ovule ratio is negatively correlated with pollen transfer efficiency in a natural community. Plant Biology 16: 843–847.

Harder L, Thomson JD. 1989. Evolutionary options for maximizing pollen dispersal of animal-pollinated plants. The American Naturalist 133: 323–344.

Harder LD, Barrett SCH, Cole WW. 2000. The mating consequences of sexual segregation within inflorescences of flowering plants. Proceedings of the Royal Society of London. Series B: Biological Sciences 267: 315–320.

Harder LD, Wilson WG. 1998. A clarification of pollen discounting and its joint effects with inbreeding depression on mating system evolution. American Naturalist 152: 684–695.

Harder LD. 1990. Behavioral responses by bumble bees to variation in pollen availability. Oecologia 85: 41–47.

Hargreaves AL, Harder LD, Johnson SD. 2009. Consumptive emasculation: the ecological and evolutionary consequences of pollen theft. Biological Reviews 84: 259–276.

Haslett JR. 1989. Interpreting patterns of resource utilization: randomness and selectivity in pollen feeding by adult hoverflies. Oecologia 78: 433–442.

Herrera MLG, Martinez del Rio C. 1998. Pollen digestion by New World bats: Effects of processing time and feeding habits. Ecology 79: 2828–2838.

Holloway BA. 1976. Pollen-feeding in hover-flies (Diptera: Syrphidae). New Zealand Journal of Zoology 3: 339–350.

Holsinger KE, Thomson JD. 1994. Pollen discounting in Erythronium grandiflorum: mass-action estimates from pollen transfer dynamics. The American Naturalist 144: 799–812.

Huang SQ, Takahashi Y, Dafni A. 2002. Why does the flower stalk of *Pulsatilla cernua* (Ranunculaceae) bend during anthesis?. American Journal of Botany 89: 1599–1603.

Humphreys EA, Skema C. 2023. Just add water: Rainfall-induced anther closure and color change in *Ripariosida hermaphrodita* (Malvaceae). Ecology and Evolution 13: e10219.

Johnson SA, Nicolson SW. 2001. Pollen digestion by flower-feeding Scarabaeidae: *Protea beetles* (Cetoniini) and monkey beetles (Hopliini). Journal of Insect Physiology 47: 725–733.

Johnson SD, Neal PR, Harder LD. 2005. Pollen fates and the limits on male reproductive success in an orchid population. Biological Journal of the Linnean Society 86: 175–190.

Lloyd DG. 1984. Gender allocation in outcrossing cosexual plants. In: Dirzo R, Sarukhán J, eds. Perspectives on plant population ecology. Sunderland, MA, USA: Sinauer Associates, 277–303.

Lloyd DG, Yates JMA. 1982. Intrasexual selection and the segregation of pollen and stigmas in hermaphrodite plants, exemplified by *Wahlenbergia albomarginata* (Campanulaceae). Evolution 36: 903–913.

Minnaar C, Anderson B, de Jager ML, Karron JD. 2019. Plant–pollinator interactions along the pathway to paternity. Annals of Botany 123: 225–245.

Minnaar C, Anderson B. 2019. Using quantum dots as pollen labels to track the fates of individual pollen grains. Methods in Ecology and Evolution 10: 604–614.

Minnaar C, De Jager ML, Anderson B. 2019. Intraspecific divergence in floral-tube length promotes asymmetric pollen movement and reproductive isolation. New Phytologist 224: 1160–1170.

Moir M, Anderson B. 2023. Pollen layering and male–male competition: Quantum dots demonstrate that pollen grains compete for space on pollinators. American Journal of Botany 110: e16184.

Moir M, Johson SD, Anderson B. 2022. Remarkable floral colour variation in the functionally specialized fly-pollinated iris, *Moraea lurida*. Botanical Journal of the Linnean Society 200: 218–232.

Portman ZM, Orr MC, Griswold TL. 2019. A review and updated classification of pollen gathering behavior in bees (Hymenoptera: Apoidea). Journal of Hymenoptera Research 71: 171–208.

Price MV, Waser NM. 1982. Experimental studies of pollen carryover: hummingbirds and *Ipomopsis aggregata*. Oecologia 54: 353–358.

R Core Team. 2020. R: A language and environment for statistical computing. Vienna, Austria: R Foundation for Statistical Computing.

Reynolds RJ, Westbrook MJ, Rohde AS, Cridland JM, Fenster CB, Dudash MR. 2009. Pollinator specialization and pollination syndromes of three related North American *Silene*. Ecology 90: 2077–2087.

Santana PC, Mulvaney J, Santana EM, Moir M, Anderson B. 2025. Competition for pollen deposition space on pollinators generates last-male advantage. Functional Ecology 39: 555–566.

Shibata A, Yumoto G, Shimizu H, Honjo MN, Kudoh H. 2025. Flower movement induced by weather-dependent tropism satisfies attraction and protection. Nature Communications 16: 4132.

Strickler K. 1979. Specialization and foraging efficiency of solitary bees. Ecology 60: 998–1009.

Thomson JD. 1986. Pollen transport and deposition by bumble bees in *Erythronium*: influences of floral nectar and bee grooming. The Journal of Ecology 74: 329–341.

Thorp RW. 2000. The collection of pollen by bees. Plant Systematics and Evolution 222: 211–223.

Ushimaru A, Watanabe T, Nakata K. 2007. Colored floral organs influence pollinator behavior and pollen transfer in *Commelina communis* (Commelinaceae). American Journal of Botany 94: 249–258.

Van der Pijl, L. 1960. Ecological aspects of flower evolution. I. Phyletic evolution. Evolution 14: 403–416.

Vasquez-Castro CA, Morel E, Garcia-Simpson B, Vallejo-Marín M. 2025. The fate of pollen in two morphologically contrasting buzz-pollinated *Solanum* flowers. bioRxiv 09.

Von Hase AV, Cowling RM, Ellis AG. 2006. Petal movement in cape wildflowers protects pollen from exposure to moisture. Plant Ecology 184: 75–87.

Westerkamp C, Classen-Bockhoff R. 2007. Bilabiate flowers: the ultimate response to bees?. Annals of botany 100: 361–374.

Westerkamp C. 1997. Keel blossoms: bee flowers with adaptations against bees. Flora 192: 125–132.

